# Transcriptomic characterization of MRI contrast with focus on the T1-w/T2-w ratio in the cerebral cortex

**DOI:** 10.1101/196386

**Authors:** Jacob Ritchie, Spiro Pantazatos, Leon French

## Abstract

Magnetic resonance (MR) images of the brain are of immense clinical and research utility. At the atomic and subatomic levels, the sources of MR signals are well understood. However, at the macromolecular scale, we lack a comprehensive understanding of what determines MR signal contrast. To address this gap, we used genome-wide measurements to correlate gene expression with MR signal intensity across the cortex in the Allen Human Brain Atlas. We focused on the ratio of T1-weighted and T2-weighted intensities (T1-w/T2-w) which is considered to be a useful proxy for myelin content. Positive correlations between myelin-associated genes and the ratio supported its use as a myelin marker. However, stronger correlations were observed for neurofilaments, and genes linked to the production of formaldehyde (which cross-links protein to create larger molecules). There was also an association with protein mass, with genes coding for heavier proteins expressed in regions with high T1-w/T2-w values. Oligodendrocyte gene markers were strongly correlated but this was not driven by myelin-associated genes, suggesting this signal is from non-myelinating satellite oligodendrocytes. We find the strongest support for the previous finding of high aerobic glycolysis in regions with low T1-w/T2-w ratio. Specifically, many mitochondrial genes were negatively correlated with T1-w/T2-w ratio. Genes up-regulated by pH in the brain were also highly correlated with the ratio, suggesting the pH gradient in mitochondria may explain the aerobic glycolysis association. Expression of protease subunit genes was also inversely associated with the ratio, in agreement with the protein mass correlation. While we corroborate associations with myelin and synaptic plasticity, differences in the T1-w/T2-w ratio appear to be more attributable to molecule size, satellite oligodendrocyte proportion, mitochondrial number, alkalinity, and axon caliber. Using disease-associated gene lists, we observed an enrichment of negative T1-w/T2-w ratio correlations with human immunodeficiency virus (HIV) associated genes. Expanding our analysis to the whole brain results in strong positive T1-w/T2-w associations for immune system, inflammatory disease, and microglial genes. In contrast, neuron markers and synaptic plasticity genes are negatively enriched. Lastly, our results vary little when our analysis is performed on T1-w or inverted T2-w intensities alone, possibly because the noise reduction properties of the ratio are not needed for postmortem brain scans. These results provide a molecular characterization of MR contrast that will aid interpretation of future MR studies of the brain.

## Introduction

Magnetic resonance imaging (MRI) is of immense clinical utility and has revolutionized studies of human brain structure and function (Lerch et al., 2017). T1-weighted and T2-weighted MR images are commonly used to study the structure of the brain. Contrast in a T1-weighted (T1-w) MR image is related to the time required for protons (hydrogen ions) to return to equilibrium magnetization after excitation by a radio-frequency pulse (T1 or longitudinal relaxation). T2-weighted (T2-w) values approximate the time required for the T2 or spin-spin vectors in the magnetic transverse plane to return to equilibrium (loss of phase coherence or alignment). Contrast in these images is associated with molecule size, iron content, diffusion/flow, pH, water content, water binding, and proton density (Koenig, 1995; MacKay and Laule, 2016; Stüber et al., 2014; Vymazal et al., 1995). *E.g.* water appears bright in T1-w imaging and dark in T2-w imaging, while the white matter of the brain is intense in T1-w images and dark gray in T2-w images. The T1-w and T2-w MRI signals have been associated with histologically assayed myelin content in a study of diseased and normal white matter (Schmierer et al., 2008). These contrasts allow identification of pathologies and segmentation of tissues, for example, white matter lesions that mark demyelination and inflammation are bright in T2-w images and black in T1-w images (Sahraian et al., 2010). While coarse tissue differences are visible across the brain and attributable to general properties, a more detailed understanding of the molecular determinants of MR image contrast in homogeneous brain regions is lacking.

To enhance contrast in the cortex, Glasser and Van Essen proposed and evaluated the ratio of the T1-w and T2-w intensities as a ‘myelin map’ of the cortex (Glasser and Van Essen, 2011). The theoretical justification for the use of the T1-w/T2-w ratio as an indicator of myelin content is that this measure attenuates the receiver-coil bias, which is present in both images, while increasing the contrast ratio for myelin (Glasser and Van Essen, 2011). Since the noise in the two images is largely uncorrelated, combining two independent measurements of the myelin-content signal should increase the accuracy of the measurement. By taking the T1-w/T2-w ratio across the cortical surface as representative of local relative myelin content, it is possible to use this estimate of the cortical myeloarchitecture to help generate finer cortical segmentations (Glasser et al., 2016). The T1-w/T2-w ratio was found to identify more robust markers of schizophrenia than was possible with the individual T1-w or T2-w images (Ganzetti et al., 2015; Iwatani et al., 2015). It has also been used for segmentation of the human habenula and to assay hippocampal myelin in subjects with post-traumatic stress disorder (Chao et al., 2015; Kim et al., 2016). In addition to myelin content, aerobic glycolysis and synaptic plasticity have been associated with the T1-w/T2-w ratio (Glasser et al., 2014).

This paper focuses on the Allen Human Brain Atlas (AHBA) which provides transcriptomic information for six *post mortem* brains (Hawrylycz et al., 2012). As a transcriptomic data set, the AHBA is unprecedented in its size and scope, representing a comprehensive "all genes, all structures" view of the human brain from both an anatomical and transcriptomic perspective (Shen et al., 2012). This data has been integrated with neuroimaging based data from positron emission tomography (PET) studies (Rizzo et al., 2016), fMRI activation studies (Fox et al., 2014) and connectivity (Hawrylycz et al., 2015; Richiardi et al., 2015; Romme et al., 2017). These integrative studies combined imaging data from other studies and subjects with the transcriptomic profiles of the six donor brains [reviewed in (Mahfouz et al., 2017)]. Studies have also associated expression with connectivity obtained from the diffusion tensor images of two of the Atlas brains (Forest et al., 2017; Li et al., 2016). However, to our knowledge, no studies have used the publically available MR images of the six donor brains to examine relationships between gene expression and signal intensity.

The development of single-cell RNA sequencing techniques has enabled investigation of heterogeneity among transcriptomic profiles of different classes of cells in the adult human brain (Darmanis et al., 2015), as well as in the mouse cortex (Tasic et al., 2016; Zeisel et al., 2015). This provides an additional opportunity for analysis of the degree to which locally enriched marker genes are related to the intensity of the T1-w/T2-w ratio.

The wide research and clinical use of structural neuroimaging motivates our work to characterize the molecular basis of contrast in these images. In the current paper, we investigate whether the expression of genes associated with specific diseases, cell-types, cellular components, processes, or functions have higher or lower spatial correlations with magnetic resonance intensity (Figure 1). Like the majority of neuroimaging studies, we focus on the cerebral cortex. We primarily characterize the T1-w/T2-w ratio but note that it is highly correlated with the T1-w and T2-w intensities, as expected.

**Figure 1.**
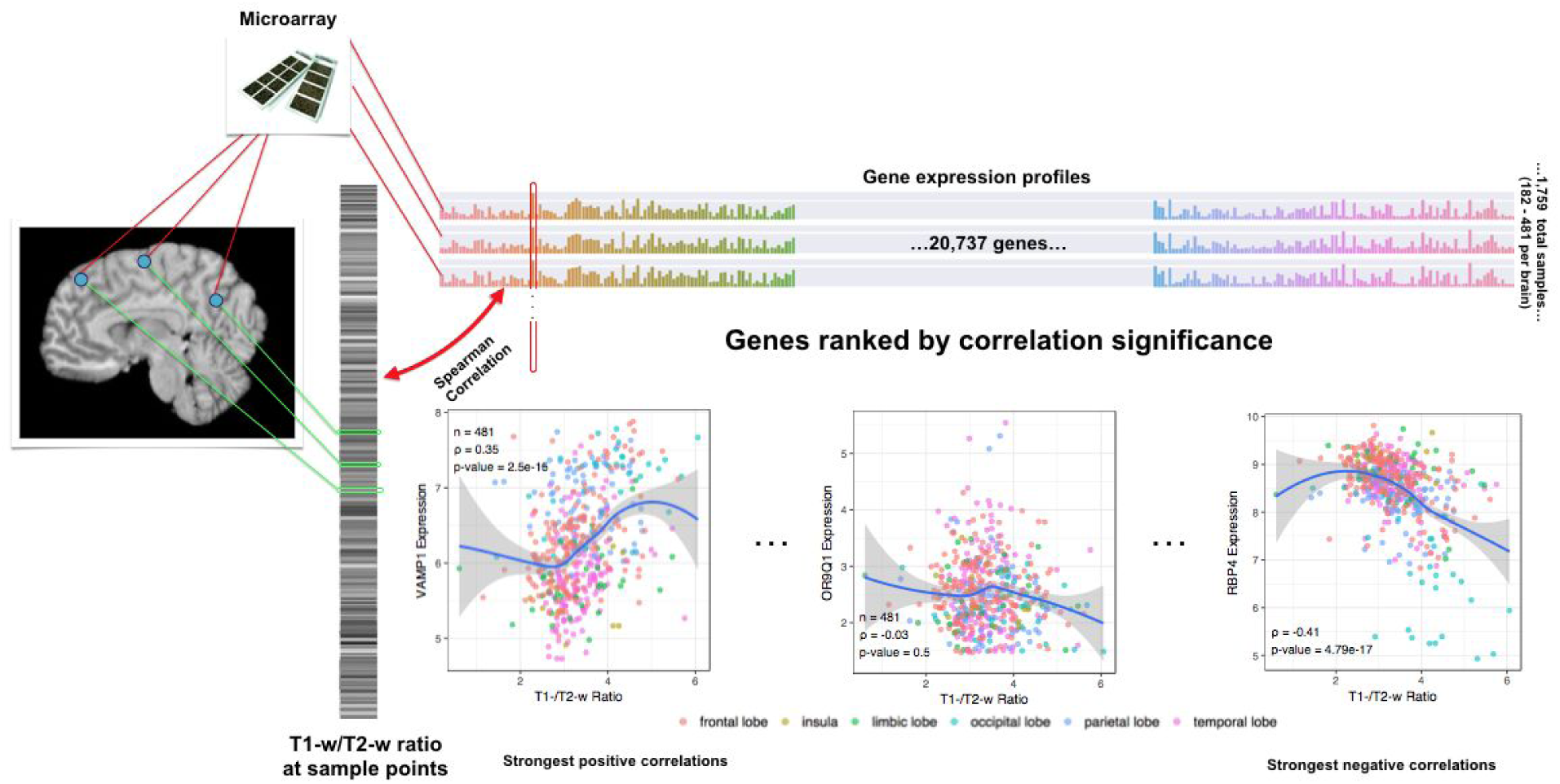
Overview of the correlation analysis. MR intensities are extracted from the images on the left, depicted as a single grayscale vector which each bar corresponding to an Allen Atlas sample. The colored bar on top visualizes the expression matrix with samples as rows and genes as columns. Each gene/column forms an expression profile across the cortex (outlined in red) and is correlated with the grayscale intensity vector.

## Materials and Methods

### Gene Expression and MRI Data

The Allen Human Brain Atlas (AHBA) is a multimodal dataset containing comprehensive transcriptomic, neuroimaging and histological information for brains obtained from 6 healthy adult human donors (1 female and 5 male, aged 24-57 years) (Hawrylycz et al., 2012). Custom 64K Agilent microarrays were used to assay genome-wide expression in 3,702 spatially-resolved samples. Full details of the procedure used by Allen Institute researchers are available in the AHBA technical white paper (http://help.brain-map.org/display/humanbrain/Documentation).

Using Python, we summarized the expression levels of the 58,692 probes to obtain average expression levels for each of the 20,737 gene transcripts. Previously provided summary statistics were used to provide measures of average expression and consistency across donors in large cortical regions (French and Paus, 2015).

In addition to transcriptomic measurements, structural postmortem MR images were acquired for the same donors by the Allen Institute on 3T Siemens Magnetom Trio scanners (Erlangen, Germany). There was variation in imaging parameters and sites across the subjects, including changes to pulse sequences, which is recorded in the AHBA technical white paper. Two brains were imaged *in cranio* and the remaining four were imaged *ex cranio*. We obtained the T1-w and T2-w MR images directly from the AHBA. Bias field correction was performed using the SPM12 software package (http://www.fil.ion.ucl.ac.uk/spm/software/spm12/).

For each transcriptomic sample, histological information allowed for a determination of the native MRI coordinates and brain region from which the sample was taken. This information was published by the AIBS along with a structural part-of ontology or reference atlas. For our main analysis, we restricted our analysis to the measurements located in the cerebral cortex (region ID of 4008 in the AHBA reference atlas) so that results would be directly applicable to characterization of the T1-w/T2-w ratio within the cortex. To focus on the neocortex, allocortical regions (hippocampal formation and piriform cortex, region IDs 4249 and 10142) from the AHBA cerebral cortex region were excluded.

In addition to the T1-w/T2-w image, the analysis was repeated for the T1-w and T2-w images, so that comparison of results could be performed between the three imaging measures.

### Correlation Analysis

Within each donor, we computed Spearman correlation between intensity of the T1-w/T2-w image at the provided native MRI coordinates and expression for each gene across the samples. Across the donors, Fisher’s method was used to generate a single combined meta p-value for each gene and direction. Then, multiple test correction was performed using the Benjamini–Hochberg false discovery rate procedure (Benjamini and Hochberg, 1995).

To obtain a genome-wide ranking of genes for enrichment analysis we ordered genes by the meta p-value and direction (from the positively correlated gene with the lowest p-value to the negatively correlated gene with the lowest p-value).

### Gene Ontology Enrichment Analysis

A Gene Ontology (GO) enrichment analysis was performed on the genome-wide gene rankings generated from the signed p-values. Annotations for GO gene groups were taken from the GO.db and org.Hs.eg.db packages in R (Carlson, 2017a, 2017b). Annotations were dated March 29, 2017. We calculated the area under the receiver operator curve (AUROC) statistic for GO groups that were over 10 and less than 200 genes, after intersection with genes present in the Allen microarray data (6885 GO groups annotating 13384 unique genes). The AUROC for a set of genes is equivalent to the probability that a gene associated with that set will be found first in the genome-wide ranking compared to an unassociated gene. Because the ranking arranges genes from strongest positive correlation to strongest negative correlation, an AUROC > 0.5 for a given group implies that samples with high expression of this gene group will be more likely to have a brighter T1-w/T2-w value. AUROC values were generated using the tmod transcriptomics enrichment analysis package in R (Weiner and Domaszewska, 2016). The Mann–Whitney U test was used to determine statistical significance. Multiple test correction of the resultant p-values for the many GO groups tested was performed using the Benjamini–Hochberg false discovery rate procedure.

We investigated eight myelin-related GO groups, which were selected by filtering for GO terms containing the strings “myelin” or “ensheathment” and excluding the strings “peripheral” and “sphingomyelin”. Because of the *a priori* hypothesis of a correlation between myelin-associated genes and the T1-w/T2-w ratio, we performed a separate analysis for these GO groups of interest and did not consider non-myelin-related GO groups while performing multiple test correction.

### Myelin Fraction Marker Genes

A transcriptomic study of myelin extracted from whole brain provided a list of genes that were more abundant in myelin from six-month-old mice compared to cortex (Thakurela et al., 2016). This list was obtained from Supplement Table S2 of Thakurela et al., which contains genes with a normalized read count of greater than 100 which are at least twice as abundant in myelin than cerebral cortex.

### Cell type enriched genes

Lists of genes that mark specific transcriptomic cell-types in the mouse and human brain were used as a source of cell type markers. Enrichment of these gene lists was determined using the previously described AUROC statistic. All cell type-enriched gene lists taken from mouse studies were first mapped to human homologs using the Homologene database (NCBI Resource Coordinators, 2016) and Ogan Mancarci’s “homologene” R package (https://github.com/oganm/homologene).

### Zeisel Marker Genes

Cell type-specific genes from the mouse brain were obtained from a single cell analysis of 3,005 cells from the adult mouse somatosensory cortex (S1) and hippocampus (Zeisel et al., 2015). We used genes from the nine modules provided in Supplemental Table 1 of Zeisel et. al. (2015). These modules were identified by clustering of both cells and genes and correspond to 9 major classes (interneurons, SS pyramidal neurons, CA1 pyramidal neurons, oligodendrocytes, microglia, endothelial cells, mural cells, astrocytes and ependymal cells).

### Neuroexpresso Marker Genes

We used cell type marker lists obtained from a cross-laboratory analysis of cell type specific transcriptomes (Ogan Mancarci et al., 2017). Ogan et. al. aggregated publicly-available gene expression values from microarray and single-cell RNA-seq studies to identify marker genes for 36 major cell types. We only used the markers obtained from cortex datasets, which included marker lists for astrocytes, endothelial cells, oligodendrocytes, OPCs, pyramidal neurons, GABAergic neurons (further labelled as PV, RelnCalb and VIPReln), and microglia (further classified as activation state independent, activated, and deactivated). The gene lists were obtained from https://github.com/oganm/neuroExpressoAnalysis.

### Darmanis Marker Genes

From the first single cell transcriptome analysis of healthy human cortex, we used marker gene lists corresponding to six main cell types (Darmanis et al., 2015). This study profiled expression in 466 cells from the human cortical tissue of eight adult subjects and four embryos. Clustering was employed to group the cells into six main groups (astrocytes, oligodendrocytes, oligodendrocyte precursor cells (OPCs), neurons, microglia, and endothelial cells) based on their transcriptomic profile, which was obtained using RNA sequencing (Darmanis et al., 2015). We used the top 21 enriched genes in each of the unbiased groups from Table S3 of Darmanis et. al. and excluded the gene groups associated with fetal cell types.

### Zeng Marker Genes

Cortical layer enriched genes were obtained from an *in-situ* hybridization (ISH) analysis performed by (Zeng et al., 2012)). In this study, 996 genes assayed in the human visual cortex and temporal cortex. The ISH images were annotated for laminar and cell-type specific expression. Specifically, we used annotations in the ‘Cortical marker (human)’ column of Supplementary Table S2 (Zeng et al., 2012). The use of these marker genes for enrichment analysis was inspired by an early version of Burt et al. (2017).

### pH Associated Genes

A cross-laboratory analysis of human gene expression profiles provided a source of pH associated genes (Mistry and Pavlidis, 2010). This meta-analysis tested the effects of brain pH on gene expression in eleven datasets that primarily assayed the frontal cortex of the normal human brain. Genes up- and down-regulated with pH were obtained from Supplementary Table 10 (Mistry and Pavlidis, 2010). A subset of core pH up-regulated genes with robust associations was also used.

### Disease gene lists

Disease-associated gene sets were downloaded from the Phenocarta database on October 28th, 2016. Phenocarta consolidates data on gene-to-phenotype mappings taken from 17 distinct resources (Portales-Casamar et al., 2013). The three sources that provide the majority of the annotations in the database are the Disease and Gene Annotations, The Rat Genome Database and the Comparative Toxicogenomics Database (Davis et al., 2017; Peng et al., 2013; Shimoyama et al., 2015). We investigated all disease-associated gene lists in the size range of 5-200 genes, a total of 1,177 diseases.

### Availability

Scripts, supplemental tables, and data files for reproducing the analyses are available online at http://www.github.com/jritch/mri_transcriptomics and https://figshare.com/articles/Transcriptomic_characterization_of_MRI_contrast_focused_on_the_T1-w_T2-w_ratio_in_the_cerebral_cortex/5270926.

Genes are then sorted from strong positive to strong negative correlation for gene set enrichment analysis to test if a particular set is enriched for positive or negative correlations. *VAMP1* (highest positive correlation), *OR9Q1* (no correlation), and *RBP4* (strong negative correlation) provide examples across the correlation range using data from donor H0351.2001.

## Results

Allen Human Brain Atlas samples per donor ranged from 182 to 481 cortical regions. Justifying our focus on T1-w/T2-w ratio, we found intensities of T1-w, T2-w and their ratio within the single brains were highly correlated across the samples (absolute Spearman’s rho > 0.79). In single brains, correlations between gene expression and T1-w/T2-w ratio reach Spearman’s correlation values ranging from −0.47 to 0.41. Combined, median correlation across the six donors ranged from −0.29 to 0.26. After multiple test correction, 1,081 positively correlated genes and 2,499 negatively correlated were identified at p_FDR_ < 0.05. Inconsistency across brains is observed with only 45% of the 3,580 significant genes showing the same correlation direction across all six brains. In the top ten positively correlated genes we note two calcium associated genes (*SYT2* and *STAC2*), and two sodium channel subunits (*SCN1B* and *SCN4B*) (genome-wide list in Supplement Table 1). We also note Estrogen Related Receptor Gamma (*ESRRG*), which is ranked 4th. A large genetic study found *ESRRG* to have a highly suggestive association with brain infarcts (defined by abnormal MRI signal intensity) (Debette et al., 2010). While no myelin-associated genes are in the top ten, *AGPAT9*/*GPAT3* which is involved in the production of lysophosphatidic acid ranks 3rd. Lysophosphatidic acid is associated with multiple sclerosis (Schmitz et al., 2017) and its receptor regulates myelination in the mouse cerebral cortex (García-Díaz et al., 2015). In the ten most negatively correlated genes we note that Retinol Binding Protein 4 (*RBP4*) is ranked second, its correlation in six brains is shown in Figure 2. Like *GPAT3*, we note that *RBP4* is also indirectly associated with multiple sclerosis. Specifically, retinol levels in blood were correlated with new lesions in multiple sclerosis patients, as detected with MR imaging (Løken-Amsrud et al., 2013).

**Figure 2.**
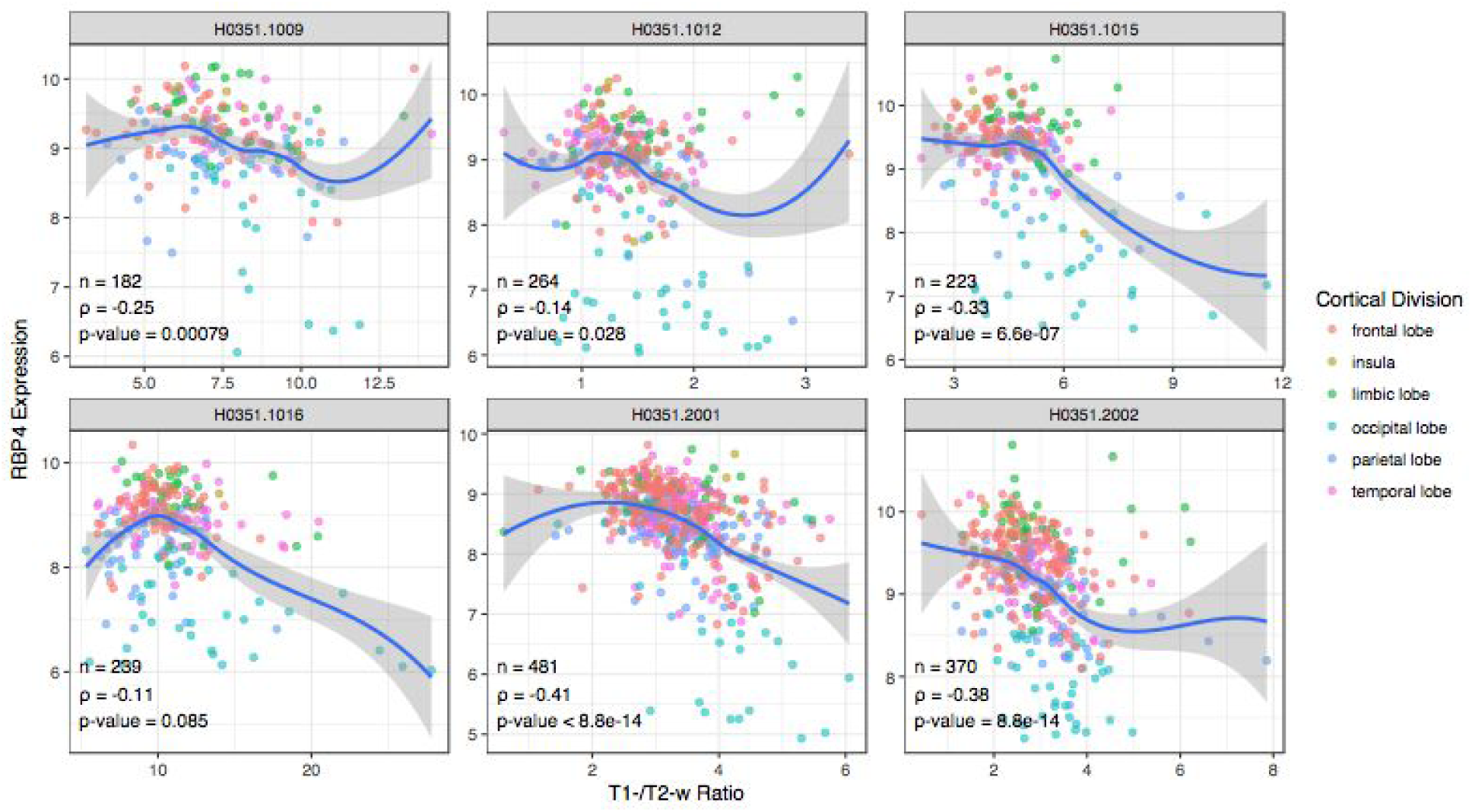
Sample-wise plots of T1-w/T2-w ratio and RBP4 gene expression. Each donor represented as a single scatter plot of samples. Local polynomial regression curves are plotted in blue. Samples are coloured by their cortical lobe/division.

### Gene Ontology Enrichment Analysis

Beyond the top ten lists, we employed gene set enrichment analysis to summarize the genome-wide results that contain a large number of significantly correlated genes.

Of the 6,443 GO groups considered in the enrichment analysis, 273 were significantly enriched (p_FDR_<0.05). Like the genewise results there more negative than positive relations with 225 GO groups negatively enriched (AUROC < 0.5) and 48 positively enriched (AUROC > 0.5).

The top ten most enriched GO groups are related to mitochondrial and peptidases (Table 1). ROC curves and raster plots are provided for select groups in Figure 3 with full results in Supplement Table 2. The “peptidase complex“, “regulation of cellular amino acid metabolic process“ and “endopeptidase complex“ GO groups overlap and primarily contain of proteasome subunits. Mitochondrial protein complex is the top-ranked group (127 genes, AUROC = 0.3) and seven of the ten top groups contain similar genes. For example, the “ribosomal subunit” genes are primarily specific to mitochondrial ribosomes (the top-ranked gene for this group is mitochondrial ribosomal protein L28). Mitochondria maintain a pH gradient by pumping hydrogen protons from the mitochondrial matrix (pH = 7.7) to the inner mitochondrial space (pH = 6.8, more acidic than the cytosol), supporting oxidative phosphorylation (Santo-Domingo and Demaurex, 2012). To more directly test pH associated genes, we used genes up- and down-regulated with pH in postmortem human brain samples (Mistry and Pavlidis, 2010). In agreement with the mitochondrial enrichment, we found that pH up-regulated genes are negatively correlated with T1-w/T2-w ratio (1109 genes, AUROC=0.324, p < 0.0001) with a core set of 13 genes showing stronger enrichment (AUROC = 0.17, p < 0.0005) and the pH down-regulated genes showing the opposite enrichment (AUROC = 0.546, 2,572 genes, p < 0.0005).

**Table 1:** Top 10 Gene Ontology groups for T1-w/T2-w ratio in cortex with negative enrichment (AUROC < 0.5). CC: cellular component; MF: molecular function; BP: biological process.

| Name | Gene Count | AUROC | p<sub>FDR</sub> | Aspect | AUROC .T1 | AUROC .T2 |
| --- | --- | --- | --- | --- | --- | --- |
| mitochondrial protein complex | 127 | 0.309 | 1.97E-10 | CC | 0.331 | 0.644 |
| structural constituent of ribosome | 140 | 0.318 | 1.97E-10 | MF | 0.324 | 0.635 |
| ribosomal subunit | 162 | 0.339 | 1.7E-09 | CC | 0.351 | 0.624 |
| inner mitochondrial membrane protein complex | 106 | 0.303 | 2.17E-09 | CC | 0.316 | 0.647 |
| mitochondrial membrane part | 166 | 0.35 | 2.03E-08 | CC | 0.371 | 0.617 |
| peptidase complex | 80 | 0.29 | 5.32E-08 | CC | 0.319 | 0.683 |
| regulation of cellular amino acid metabolic process | 59 | 0.269 | 0.00000039 | BP | 0.295 | 0.711 |
| large ribosomal subunit | 103 | 0.325 | 0.00000039 | CC | 0.341 | 0.629 |
| endopeptidase complex | 65 | 0.282 | 0.00000047 | CC | 0.311 | 0.692 |
| oxidative phosphorylation | 95 | 0.322 | 0.00000066 | BP | 0.342 | 0.628 |

**Figure 3:**
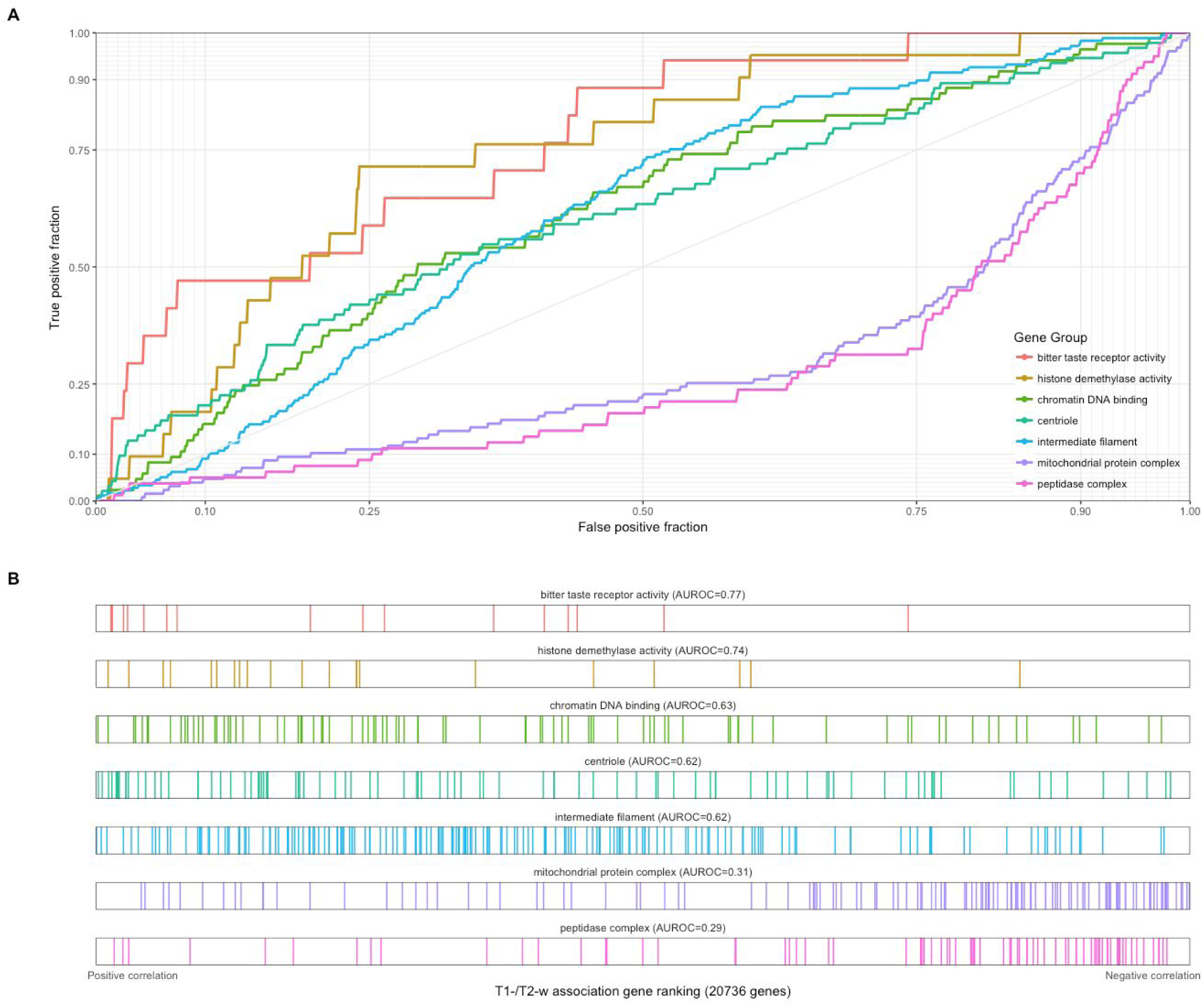
GO groups with positive and negative highest associations with T1-w/T2-w ratio intensities in the cortex. (A) ROC curves for GO groups selected from Tables 1 and 2. The curves show the proportion of GO group genes that overlap (y-axis, true positive fraction) in varying lengths of the T1-w/T2-w gene ranking (approximated by the x-axis, false positive fraction). Colored lines mark genes in different GO groups. (B) Distributions of the seven significant GO groups across the T1-w/T2-w associated gene ranking with each annotated gene representing a single coloured line. The GO groups are arranged from most positively (bitter taste receptor activity) to most negatively enriched (peptidase complex).

**Table 2:** Top 10 Gene Ontology groups for T1-w/T2-w ratio in cortex with positive enrichment (AUROC > 0.5). CC: cellular component; MF: molecular function; BP: biological process.

| Name | Gene Count | AUROC | $p_{FDR}$ | Aspect | AUROC. T1 | AUROC. T2 |
| --- | --- | --- | --- | --- | --- | --- |
| intermediate filament | 177 | 0.62 | 0.0000050 | CC | 0.598 | 0.39 |
| keratin filament | 85 | 0.655 | 0.000069 | CC | 0.628 | 0.365 |
| chromatin DNA binding | 85 | 0.626 | 0.0019 | MF | 0.6 | 0.386 |
| centriole | 93 | 0.621 | 0.0019 | CC | 0.626 | 0.379 |
| transcriptional repressor activity, RNA polymerase II transcription regulatory region sequence-specific binding | 177 | 0.586 | 0.0025 | MF | 0.58 | 0.413 |
| bitter taste receptor activity | 17 | 0.774 | 0.0030 | MF | 0.765 | 0.209 |
| histone demethylase activity | 21 | 0.744 | 0.0034 | MF | 0.743 | 0.262 |
| basal part of cell | 45 | 0.664 | 0.0043 | CC | 0.681 | 0.362 |
| histone demethylation | 23 | 0.726 | 0.0050 | BP | 0.724 | 0.277 |
| core promoter binding | 157 | 0.585 | 0.0069 | MF | 0.568 | 0.431 |

Although the positively enriched GO groups have weaker enrichment, they are more diverse (Table 2 and Figure 3). Intermediate filament is the top-ranked positively enriched group and its most correlated genes are the brain-associated *NEFH*, *NEFM*, *INA*, *NEFL*, and *DST*. The neurofilament genes are ranked in order of decreasing weight, suggesting a relationship with axon caliber. The remainder of the genes in this group are keratin or keratin-associated proteins. Keratin genes have little or no expression and the brain and are not consistently expressed across the 6 brains (French and Paus, 2015), suggesting this result is driven primarily by enrichment of the neurofilaments genes. The second-ranked ‘keratin filament’ group contains only these keratin genes and may be a false positive, as it contains only one gene that is significantly correlated with T1-w/T2-w ratio (after correction, positive correlation). The “histone demethylase activity” and “histone demethylation” groups also contain a large number of overlapping genes which are mostly lysine demethylases. In Table 1 and 2, the T1-w/T2-w ratio AUROC values are slightly more extreme than T1-w or T2-w results, suggesting a better signal.

We next performed a targeted analysis of myelin-related gene ontology groups. Three of the eight myelin groups (“ensheathment of neurons”, “myelination” and “myelin assembly”) were found to be positively enriched with the T1-w/T2-w ratio (p_FDR_ < 0.05, Table 3 and Figure 4). However, we note that these GO groups strongly overlap and that the majority were positively enriched. Relative to all tested GO groups, the myelin-related groups are less enriched than over 200 other GO groups. Also, they are ranked slightly higher in the enrichment results in separate T1-w and T2-w analyses (AUROC values are similar). An independent list of myelin marker genes obtained from transcriptomic profiling of whole mouse brain myelin fraction has similar enrichment (1618 genes, AUROC = 0.58, p_FDR_<0.000001) (Thakurela et al., 2016). The second process of interest is synaptic plasticity, which has been previously associated with the T1-w/T2-w ratio (Glasser et al., 2014). The “regulation of synaptic plasticity” GO group is negatively enriched (AUROC=0.40, p_FDR_=0.005, rank=106), as is the”long-term synaptic potentiation” GO group (AUROC=0.38, p_FDR_=0.02, rank=189).

**Figure 4:**
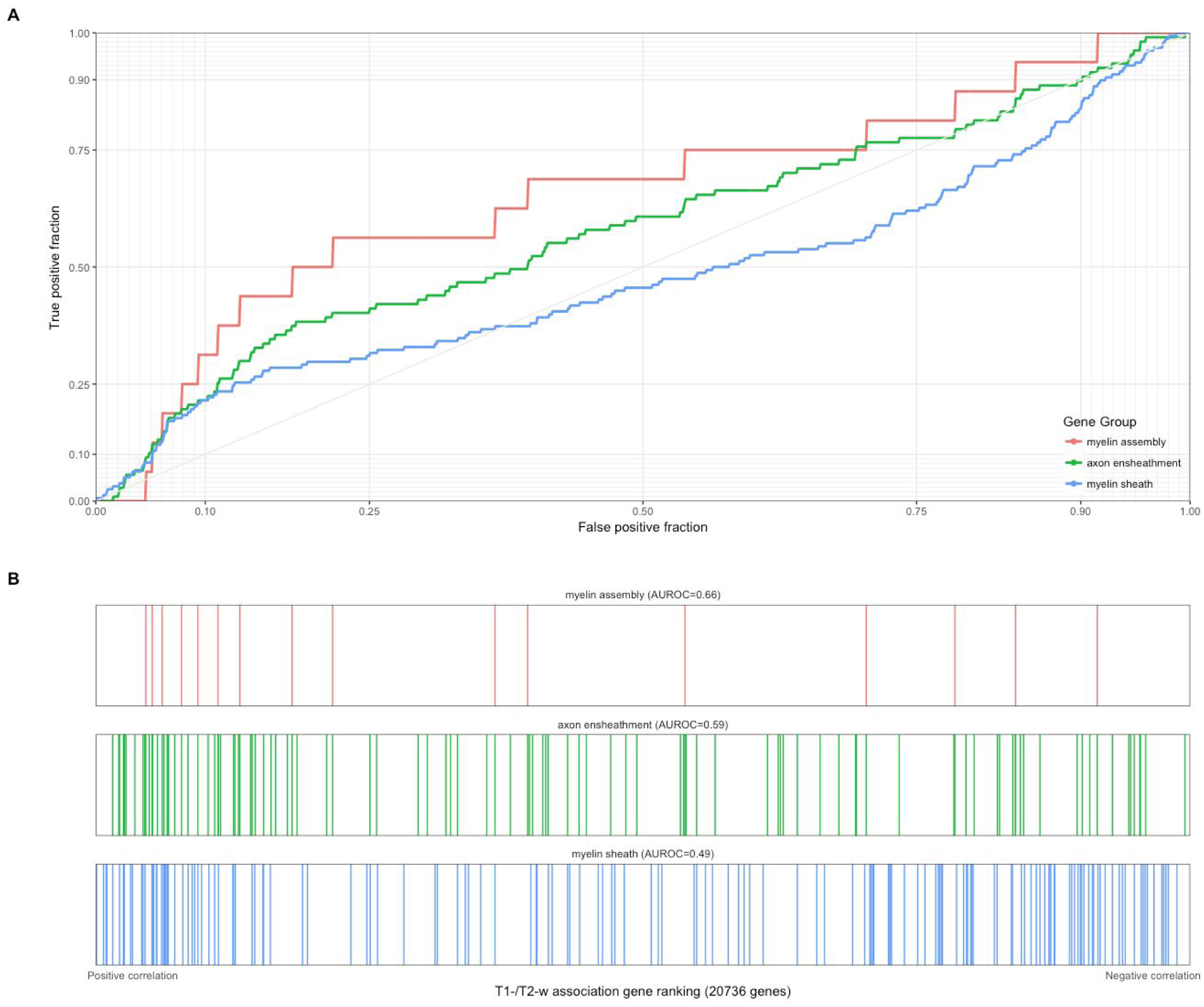
Associations with T1-w/T2-w ratio intensities for selected myelin-related GO groups. (A) ROC curves for GO groups selected from Table 3. The curves show the proportion of GO group genes that overlap (y-axis, true positive fraction) in varying lengths of the T1-w/T2-w gene ranking (approximated by the x-axis, false positive fraction). Colored lines mark genes in different GO groups. (B) Distributions of the three significant GO groups across the T1-w/T2-w associated gene ranking with each annotated gene representing a single coloured line.

**Table 3:**
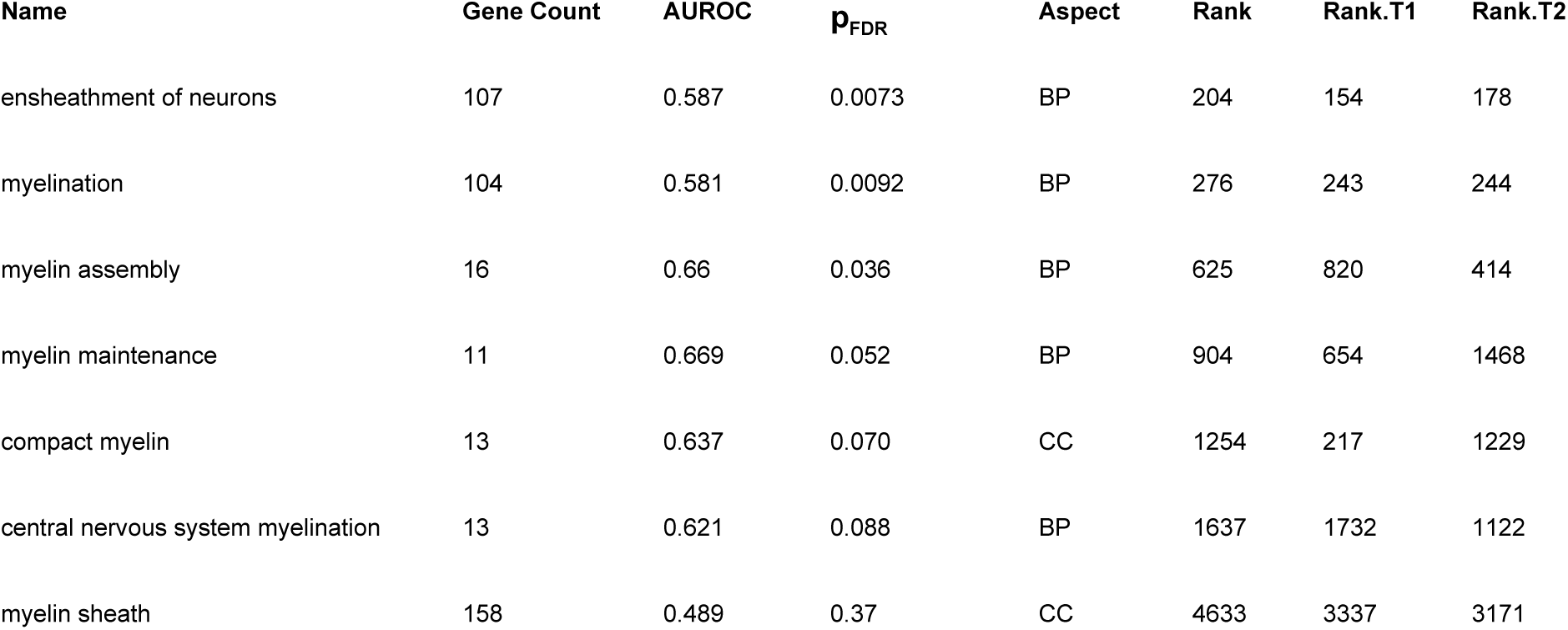
Enrichment statistics for myelin-related Gene Ontology groups using the T1-w/T2-w ratio ranking.

| Name | Gene Count | AUROC | $p_{FDR}$ | Aspect | Rank | Rank.T1 | Rank.T2 |
| --- | --- | --- | --- | --- | --- | --- | --- |
| ensheathment of neurons | 107 | 0.587 | 0.0073 | BP | 204 | 154 | 178 |
| myelination | 104 | 0.581 | 0.0092 | BP | 276 | 243 | 244 |
| myelin assembly | 16 | 0.66 | 0.036 | BP | 625 | 820 | 414 |
| myelin maintenance | 11 | 0.669 | 0.052 | BP | 904 | 654 | 1468 |
| compact myelin | 13 | 0.637 | 0.070 | CC | 1254 | 217 | 1229 |
| central nervous system myelination | 13 | 0.621 | 0.088 | BP | 1637 | 1732 | 1122 |
| myelin sheath | 158 | 0.489 | 0.37 | CC | 4633 | 3337 | 3171 |

### Cell Type Marker Enrichment Analysis

We first tested if the Zeisel marker genes were enriched in the T1-w/T2-w associations (Table 4, Figure 5). These marker genes were obtained from a single cell analysis of the adult mouse somatosensory cortex (S1) and hippocampus. Oligodendrocyte genes are the most strongly enriched with high positive correlations with T1-w/T2-w intensities. Pyramidal neurons from the hippocampus are negatively enriched (AUROC = 0.39, p_FDR_ < 0.0001), while those from S1 are not significantly enriched. Microglia, interneuron, and astrocyte marker genes are also negatively enriched in the T1-w/T2-w correlation ranking. To test robustness we used a second set of markers from the Neuroexpresso cross-laboratory database (Ogan Mancarci et al., 2017). Similar results were obtained, with enrichment for oligodendrocyte, pyramidal, microglia (markers of the inactivated state), and astrocyte markers (Supplemental Table 3). In contrast to the Zeisel interneuron finding, none of the three Neuroexpresso GABAergic subtypes are enriched. Also, the Neuroexpresso but not Zeisel endothelial markers are positively enriched (AUROC = 0.57, p_FDR_ < 0.02).

**Table 4:** Enrichment statistics for Zeisel marker gene sets using the T1-w/T2-w ratio ranking.

| **Cell-type or class** | **Gene Count** | **AUROC** | **$p_{\text{FDR}}$** |
| --- | --- | --- | --- |
| Oligodendrocyte | 368 | 0.621 | 8.0E-15 |
| Neuron CA1 pyramidal | 327 | 0.393 | 1.4E-10 |
| Microglia | 355 | 0.407 | 5.3E-09 |
| Interneuron | 298 | 0.437 | 0.00060 |
| Astrocyte | 203 | 0.439 | 0.0070 |
| Mural | 123 | 0.553 | 0.086 |
| Neuron S1 pyramidal | 220 | 0.461 | 0.086 |
| Ependymal | 370 | 0.477 | 0.12 |
| Endothelial | 296 | 0.519 | 0.12 |

**Figure 5.**
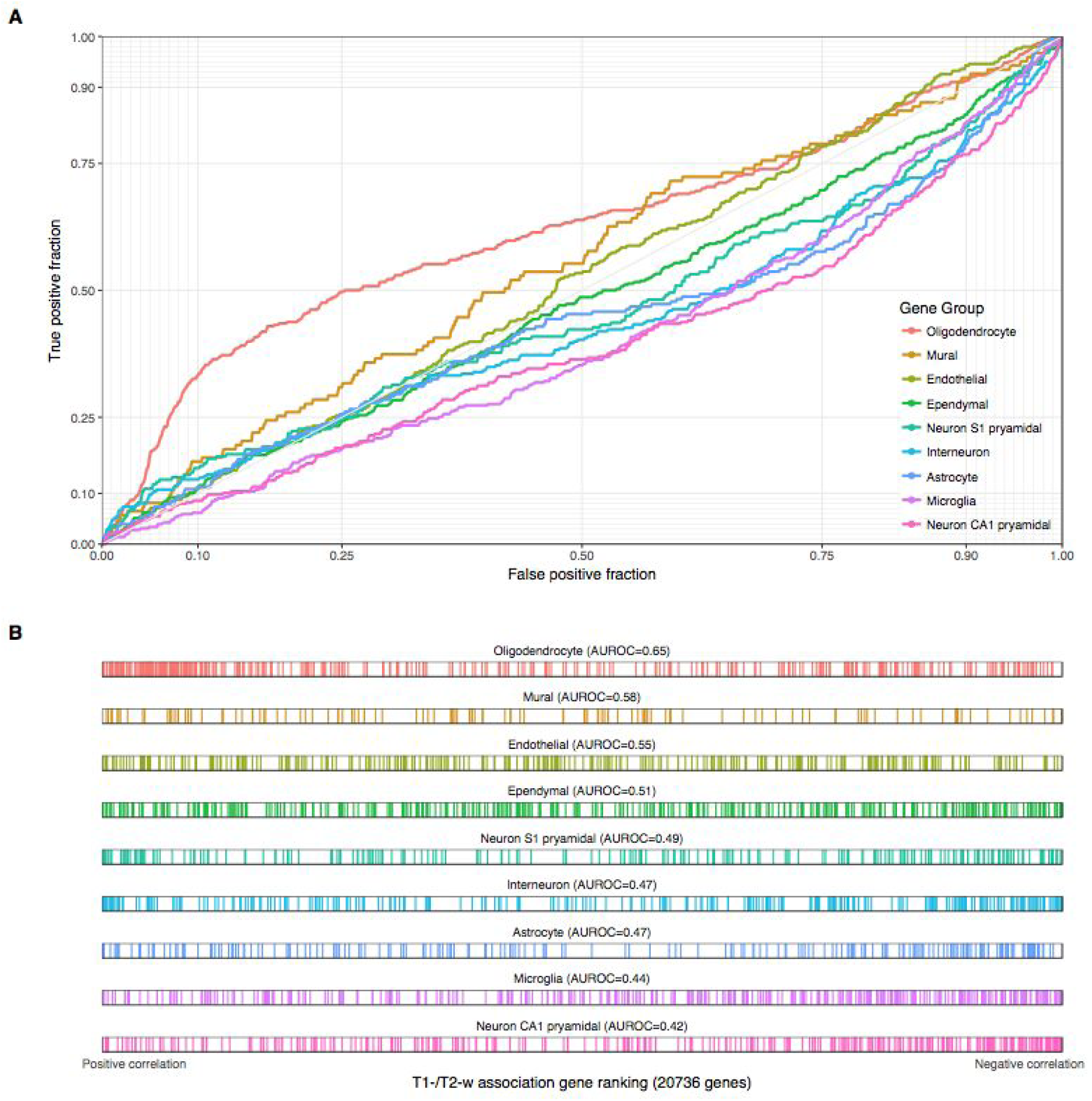
Associations with T1-w/T2-w ratio intensities for Zeisel cell type marker genes. (A) ROC curves showing the proportion of marker genes that overlap (y-axis, true positive fraction) in varying lengths of the T1-w/T2-w gene ranking (approximated by the x-axis, false positive fraction). Colored curves mark genes in different cell type marker lists. (B) Distributions of the cell type markers across the T1-w/T2-w associated gene ranking. Each marker gene is represented by a single coloured line.

**Figure 6.**
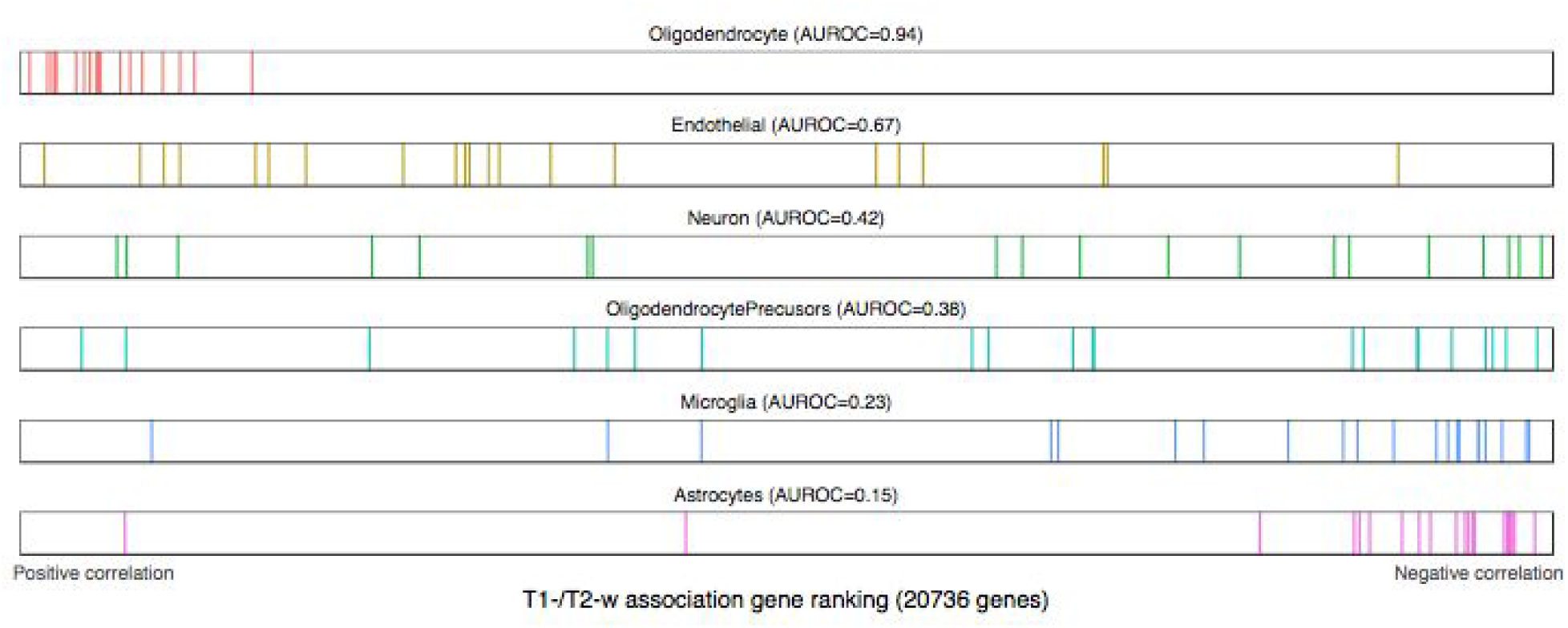
Distributions of the Darmanis cell type markers across the T1-w/T2-w associated gene ranking. Each marker gene is represented by a single coloured line. Only the neuron marker list is not significantly enriched after test correction.

We next evaluated cell type marker enrichment with human datasets to determine if there are species specific effects. While these marker gene lists are from more limited human tissue sources, their enrichment mirrors the larger mouse dataset. Using the marker gene lists from the Darmanis et al. study of the temporal cortex, we see significant enrichment for five of the six lists (Figure 6, Supplementary Table 4). Unlike the mouse lists, neuron markers are not enriched, this is possibly due to the coarse grouping of all neurons (Ziesel and NeuroExpresso contain at least 3 neuron subclass lists). The Darmanis oligodendrocyte precursor lists was enriched (AUROC = 0.35, p_FDR_ < 0.03), but the same list from NeuroExpresso was not (AUROC = 0.53, p_FDR_ > 0.05). For endothelial markers, the Darmanis list was enriched (AUROC = 0.65, p_FDR_ < 0.03), matching the direction as the NeuroExpresso list (not significant with the Ziesel list). The top three most significant lists (oligodendrocyte, astrocyte and microglia) all match the enrichment direction of the mouse results. The second human study employed in situ hybridization (ISH) to annotate expression across the cortical layers (Zeng et al., 2012). As shown in Figure 7, astrocyte and oligodendrocyte marker genes are significantly enriched in matching directions. Genes annotated as expressed in neurons in layers 2, 3, 5, 6 or interneurons are negatively enriched (AUROC < 0.42, p_FDR_ < 0.05, Table S6). Layer 4 markers contrast the other neuron lists with an AUROC of 0.5 (Figure 7). While they generally agree, we note that the human sources of cell type markers differ in size of gene lists (6-68 genes for Zeng, 19-21 for Darmanis), cortical region assayed (visual and temporal cortices), and experimental design.

**Figure 7:**
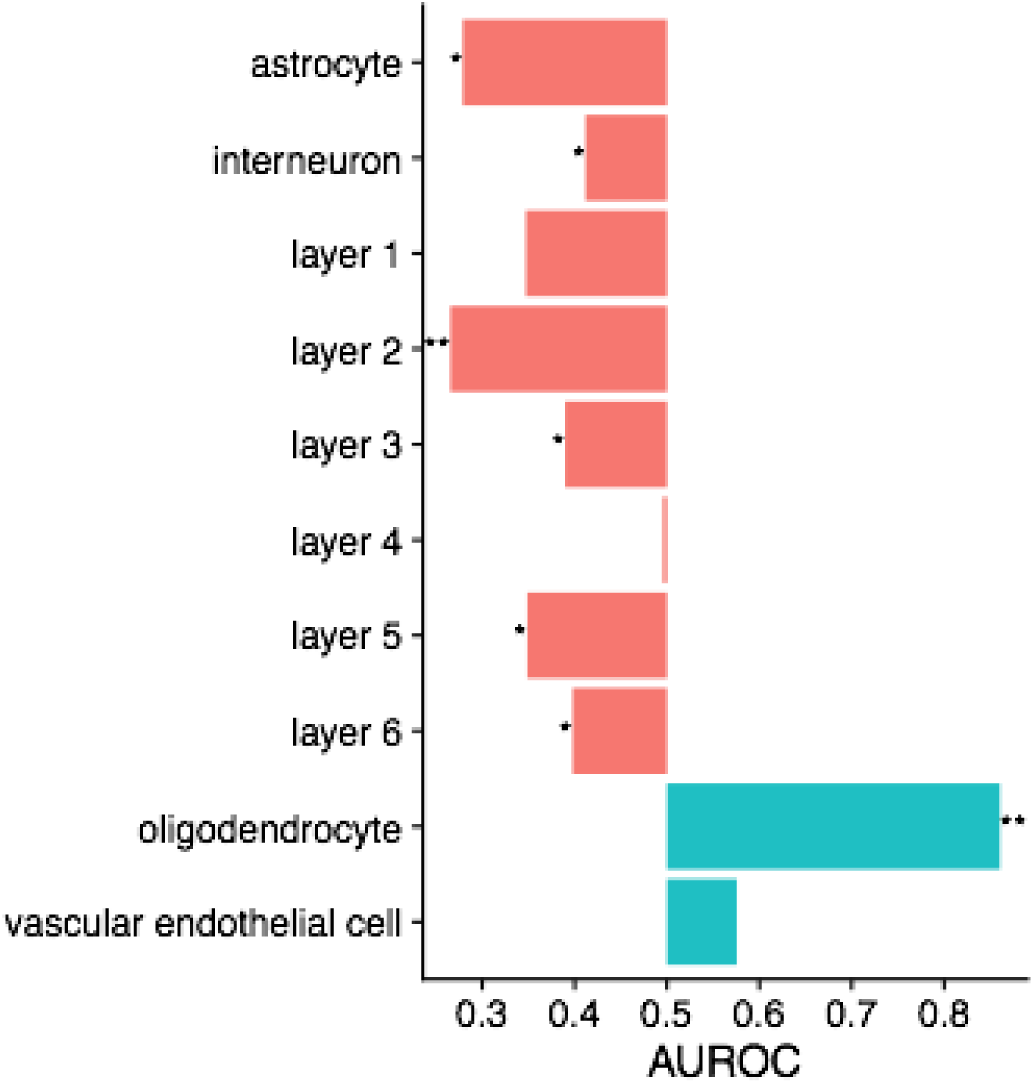
Barplot of enrichment in the T1-w/T2-w associated gene ranking across the Zeng laminar and cell-type markers. * indicates pFDR < 0.05 and ** indicates pFDR < 0.0005.

We examined the marker genes in the Darmanis oligodendrocyte markers to investigate whether the enrichment of this group was driven the by the myelin-associated genes in the list (*MAG, MOG* and *OPALIN*). These genes had lower than average correlation with T2-w/T1-w ratio intensity than the other marker genes (ranked 11, 12, and 19th of 20 genes, Supplemental Table 5). In contrast, two peptidase-coding genes, *CNDP1* and *KLK6* were top-ranked (ranked 1 and 3 of 20).The *TF* gene, which codes for transferrin an iron transport protein, was also top-ranked (4 of 20). While oligodendrocyte marker genes are strongly associated with the T1-w/T2-w ratio, it is not myelin-associated genes that are not key markers.

### Disease-associated Gene Analysis

Testing of 1,178 disease-associated gene lists revealed significant enrichment of human immunodeficiency virus (HIV) associated genes (AUROC = 0.42, p_FDR_ < 0.05). While no clear theme is evident, we note that several proteasome subunit genes are top-ranked in the HIV list. While no other disease-associated gene lists were significant after multiple correction, temporal lobe epilepsy" is ranked second (AUROC = 0.4, p_FDR_ = 0.17) and demyelinating disease is third (AUROC = 0.74, p_FDR_ = 0.17).

### Whole Brain Analysis

While we focus on the neocortex in this paper, the AHBA has comprehensively assayed the brain, allowing a wider analysis. We provide only a summary characterization of whole brain correlation results due to space constraints and because the T1-w/T2-w ratio was originally proposed for the cerebral cortex (Glasser and Van Essen, 2011).

Across the whole brain, the majority of genes are significantly correlated with T1-w/T2-w ratio intensities after multiple test correction. In a reversal of the neocortical findings, there are less negatively correlated genes (7,184) that positively correlated (10,034). The top positively enriched GO group is “MHC protein complex” (AUROC = 0.878), followed by many other immune system related groups (Supplementary Table 7). The most negatively enriched GO group is “regulation of synaptic plasticity” (AUROC = 0.30), followed by similar GO groups such as “postsynaptic density” (AUROC = 0.33). We again observe that pH up-regulated genes have higher correlations with T1-w/T2-w ratio than pH down-regulated genes. While our cell-type gene markers are from studies of the cortex, we note that endothelial and microglial markers switch from weak negative to strong positive enrichment in comparison to the neocortical results. This is observed across all three of the marker sources. We note that markers of activated microglia has the strongest enrichment of the three microglia gene lists from NeuroExpresso (AUROC = 0.71, p_FDR_ < 0.00001, Supplement Table 9). In contrast to the neocortical results, 214 of the 1178 disease-associated gene lists were significant (Supplementary Table 8). The most significant enrichment was Behcet’s disease (AUROC=0.649, p_FDR_<0.0001), an inflammatory disease that has been associated with demyelination (Al-Araji and Kidd, 2009). The most strongly negatively enriched was genes associated with Angelman’s syndrome (AUROC=0.097, p_FDR_<0.0005). Delayed or reduced myelin formation has been identified in children suffering from Angelman’s syndrome (Harting et al., 2009), as well as in mouse models of the disease (Grier et al., 2015). Most of the top listed diseases are inflammatory, positively enriched, and not clearly linked to the brain (‘pustulosis of palm and sole’ for example). The MHC protein complex GO group, microglia, endothelial cell-type, and inflammatory disease findings suggest that across the whole brain, the variance in MR signal intensity may be explained by immune or circulatory activity.

## Discussion

MR images of the brain are of immense clinical and research value. These grayscale images provide contrast at the submillimetre scale, allowing segmentation of many structures and identification of abnormalities. At the atomic and subatomic levels, the sources of MR signals are well understood. However, at the macromolecular scale, we lack a comprehensive understanding of what determines MR signal intensity. To address this gap, we have used genome-wide measurements to correlate gene expression with MR signal intensity. We found biological processes, cellular components, molecular functions, diseases, and cell-types that are related to contrast by leveraging gene annotations. This work will aid interpretation of future MR studies of the brain by providing a molecular perspective.

In this paper, we focused on the T1-w/T2-w ratio in the cerebral cortex which has been previously associated with myelin content, aerobic glycolysis and synaptic plasticity (Glasser et al., 2014; Glasser and Van Essen, 2011). These past associations were revealed by imaging (PET and histology) and not gene expression. In our results, we find the strongest support for the previous finding of high aerobic glycolysis in regions with low T1-w/T2-w ratio. Specifically, our strongest GO analysis finding is negative enrichment of mitochondrial GO groups, with high expression of mitochondrial genes in cortical regions with low T1-w/T2-w ratio. Mitochondria are the location of aerobic glycolysis and cellular respiration (via oxidative phosphorylation). Most of the genes that are anticorrelated with T1-w/T2-w ratio are located on the inner mitochondrial membrane, where a pH gradient is maintained. Crossing this membrane changes pH from 6.8 to 7.7 (Santo-Domingo and Demaurex, 2012), which is used to spin the ATP synthase complex through the transfer of hydrogen ions (H^+^). Because MR scanners target signals from H^+^, we tested genes associated with pH in postmortem brain studies (Mistry and Pavlidis, 2010). This provided a more direct comparison of H^+^ relations than aerobic glycolysis or mitochondrial genes. In agreement, we found that genes correlated with pH were also correlated with T1-w/T2-w ratio intensity. This suggests that regions with low T1-w/T2-w ratio intensities have elevated concentrations of H^+^ due to a higher density of mitochondria that are supplying energy through aerobic glycolysis.

While a clear mitochondrial-associated signal is inversely correlated with T1-w/T2-w ratio, we observe much weaker signals in the positive direction. At the global level, fewer genes and GO groups are significantly associated with T1-w/T2-w ratio in the positive direction. Also, the genes that are significantly positively correlated have lower average expression (6.34) than the negative direction (7.07, p < 0.00001). This may be caused by “transcriptional repressor activity, RNA polymerase II transcription regulatory region sequence-specific binding”, as this GO group is the 5th most significant group in the positive direction. Several top-ranked GO groups lack a clear connection to T1-w/T2-w ratio. For example, the keratin filament GO group is ranked second and contains keratin genes that are not widely expressed in the brain. This GO group has been found to be not consistently expressed across the six AHBA brains (French and Paus, 2015). We also lack confidence in the “bitter taste receptor activity” GO group, which has the highest AUC value (0.77). We note there is high sequence homology across these taste receptors and nonspecific binding may explain the enrichment of the group. However, bitter taste receptor transcripts are expressed in several regions of the rat brain, and that cells containing these transcripts are functional, being capable of responding to tastants (Singh et al., 2011). This suggests that in regions with high T1-w/T2-w ratio, less mitochondria and aerobic glycolysis result in lower transcriptional activity and weaker molecular correlations.

Genes linked to molecule size show clear enrichment in our analysis. The peptidase complex GO group which consists primarily of proteasome subunit genes is negatively enriched (ranked 6th). These genes code for proteins that degrade proteins into smaller peptides or component amino acids by hydrolysis. We believe that the positively enriched “histone demethylase activity” GO group is also related to molecule size. This group consists primarily of lysine demethylation genes. A product of the lysine demethylation reaction is formaldehyde (Shi et al., 2004).

Formaldehyde is a fixative that crosslinks proteins, creating larger molecules. Fixation of brain tissue with formalin is known to produce a bright band of T1-w intensities as fixation proceeds (Schmierer et al., 2008; Yong-Hing et al., 2005). Of the 7,652 genes in our analysis and the UniProt database (The UniProt Consortium, 2017), protein mass is correlated with T1-w/T2-w association ranking (Spearman’s rho = 0.13, p < 0.0001). For the significantly T1-w/T2-w correlated genes in the positive direction the median weight is 64,914 Daltons, for the negatively correlated genes, it is 41,536 Daltons. This contrast between molecule size, peptidase activity, and demethylases that generate formaldehyde agrees with larger molecules having more restricted molecular movement that results in brighter T1-w intensities (Koenig, 1995; Sprawls, 2000).

Genes specifically expressed in oligodendrocytes are highly expressed in regions with high T1-w/T2-w ratio. This was the strongest and most consistent cell-type in our marker enrichment analysis. In regards to transcriptional activity, we also note that oligodendrocytes have half the number of detected RNA molecules per cell when compared to neurons (Zeisel et al., 2015), in agreement with the above findings of lowly expression genes, and transcriptional repression.

Generation of the myelin sheath is considered the primary role of oligodendrocytes. However, the oligodendrocyte marker genes with the highest T1-w/T2-w correlation are not myelin-associated genes. Furthermore, while some myelin-associated GO groups are positively enriched in the T1-w/T2-w gene ranking, there are over 200 GO groups with stronger positive enrichment (Table 3). This lack of a strong myelin signal suggests that the oligodendrocyte signal we observe is driven by satellite perineuronal oligodendrocytes that are primarily non-myelinating (Battefeld et al., 2016; Takasaki et al., 2010). We emphasize that all of the samples and cell-type marker lists used were derived from cortical gray matter samples which are known to have low myelin content (MacKay and Laule, 2016). Previous association of T1-w/T2-w ratio with myelin content may be due to myelin transcripts that were transcribed in white matter and transported to distal sheaths after translation into protein. In this context, we note that the ‘paranode region of axon’, which is where the myelin sheath ends, has the highest AUROC value (0.8, 10 genes, pFDR < 0.02). However, we note that myelin basic protein is translated locally at the axon contact site (Müller et al., 2013).

Our findings suggest a positive correlation between T1-w/T2-w intensity and axon caliber. The three neurofilament genes are positively correlated with the ratio in order of decreasing weight with the heavy chain (*NEFH*) ranked 13th genome-wide. Expression of *NEFH* plays a major role in the development of high-caliber axons (Elder et al., 1998). A positive correlation has been observed between axon caliber and myelin thickness in neurons, both in developing nerves (Friede, 1972) and in the adult brain (Fraher and Dockery, 1998). *NEFH* has been linked to an oligodendrocyte associated expression pattern that is expressed at higher levels in ‘relay-like’ regions with few connections (French et al., 2011). Parvalbumin (*PVALB*), another gene in the oligodendrocyte associated expression pattern is highly ranked (86th most positive correlation).

*PVALB* marks fast-spiking GABAergic interneurons that are frequently myelinated and may explain a large proportion of myelin in the cortex (Stedehouder et al., 2017). Therefore, our finding of T1-w/T2-w ratio correlation for neurofilaments and parvalbumin provides indirect support for an association with increased myelin content.

Beyond the consistent enrichment of oligodendrocyte markers, we observe enrichment of genes that are specifically expressed in astrocyte, neuron, endothelial and microglial cells. Astrocyte and microglia marker gene sets were significantly negatively correlated with T1-w/T2-w ratio on average in all four sources of cell type markers (three sources for microglia). Endothelial marker sets had positive AUC values in all four sources but the relationship was significant in only two. Neuron markers were negatively enriched. There is not a clear difference between inter- and pyramidal neuron markers. Markers for CA1 pyramidal neurons are more strongly enriched than those from S1. While CA1 wasn’t used in our study, the CA1 markers may gauge synaptic plasticity (Shin et al., 2017). Synaptic plasticity GO groups were negatively enriched at levels similar to the neuron markers (AUROC ~= 0.4), supporting previously described links with plasticity (Glasser et al., 2014).

A negative enrichment of HIV linked genes resulted from our analysis of disease-associated genes. These correlated HIV genes include proteasome subunits, which have been previously been associated with HIV (Krishnan and Zeichner, 2004; Schubert et al., 2000; Seeger et al., 1997). In some patients, HIV RNA is detected in cerebrospinal fluid and has been associated with dementia and diffuse white matter signal abnormalities (Kugathasan et al., 2017). Beyond HIV, no other disease gene lists are significant after correction. However, we note that demyelinating disease is ranked third and is positively enriched. Also, a few top-ranked genes are indirectly related to multiple sclerosis.

In our analysis of the whole brain, we observe that immune-related genes, oligodendrocyte, activated microglia markers are positively correlated with T1-w/T2-w ratio. In contrast, neuron markers and related GO groups such as ‘postsynaptic density’ are negatively enriched. This is also reflected in the disease-associated gene set analysis with significant positive enrichment of inflammatory and negative enrichment of neural associated disease gene lists (Angelman syndrome and temporal lobe epilepsy for example). The Behcet’s disease gene list was the most enriched. Behcet’s disease is an autoimmune condition that manifests as vasculitis across many organs of the body, with neurological symptoms in 5-50% of cases (van der Knaap and Valk, 2011). In particular, it can cause lesions of the basal ganglia, midbrain, and pons, as well as white matter abnormalities in the cerebral hemispheres which are similar to the patterns of demyelination that occur in multiple sclerosis (van der Knaap and Valk, 2011). In comparison to the cortical analysis, the whole brain results show a broad neuron versus glia pattern with stronger correlations for microglia and immune-related genes.

It remains ambiguous whether the direct interpretation of the T1-w/T2-w ratio as a ‘myelin map’, as suggested by previous researchers is justified. Descriptions related to axon caliber, oligodendrocytes, or pH may be equally appropriate. While our transcriptomic perspective supported associations with aerobic glycolysis, future work is necessary to validate the precise relationship between myelin content and the T1-w/T2-w ratio.

We emphasize that our results vary little when our analysis is performed on T1-w or inverted T2-w intensities alone. While our enrichment values are improved when the ratio is used, it is possible the noise-attenuating properties of the ratio are of limited importance in our study because the MRI scans were acquired *post mortem*.

## Acknowledgements

This study was supported by a National Science and Engineering Research Council of Canada (NSERC) Discovery Grant to LF and a NIMH K01MH108721 to SP.

## Author contributions

Conception and design of the work: SP and LF. Analysis: JR, SP, and LF. Drafting and revising the work: JR and LF. All authors approved the manuscript.

## References

Al-Araji, A., and Kidd, D. P. (2009). Neuro-Behçet's disease: epidemiology, clinical characteristics, and management. Lancet Neurol. 8, 192–204.

Battefeld, A., Klooster, J., and Kole, M. H. P. (2016). Myelinating satellite oligodendrocytes are integrated in a glial syncytium constraining neuronal high-frequency activity. Nat. Commun. 7, 11298.

Benjamini, Y., and Hochberg, Y. (1995). Controlling the False Discovery Rate: A Practical and Powerful Approach to Multiple Testing. J. R. Stat. Soc. Series B Stat. Methodol. 57, 289–300.

Burt, J. B., Demirtas, M., Eckner, W. J., Navejar, N. M., Ji, J. L., Martin, W. J., et al. (2017). Hierarchy of transcriptomic specialization across human cortex captured by myelin map topography. bioRxiv, 199703. doi:10.1101/199703.

Carlson, M. (2017a). GO.db: A set of annotation maps describing the entire Gene Ontology. Available at: ftp://ctan.uib.no/pub/bioconductor/2.7/data/annotation/html/GO.db.html.

Carlson, M. (2017b). org.Hs.eg.db: Genome wide annotation for Human.

Chao, L. L., Tosun, D., Woodward, S. H., Kaufer, D., and Neylan, T. C. (2015). Preliminary Evidence of Increased Hippocampal Myelin Content in Veterans with Posttraumatic Stress Disorder. Front. Behav. Neurosci. 9, 333.

Darmanis, S., Sloan, S. A., Zhang, Y., Enge, M., Caneda, C., Shuer, L. M., et al. (2015). A survey of human brain transcriptome diversity at the single cell level. Proc. Natl. Acad. Sci. U. S. A. 112, 7285–7290.

Davis, A. P., Grondin, C. J., Johnson, R. J., Sciaky, D., King, B. L., McMorran, R., et al. (2017). The Comparative Toxicogenomics Database: update 2017. Nucleic Acids Res. 45, D972–D978.

Debette, S., Bis, J. C., Fornage, M., Schmidt, H., Ikram, M. A., Sigurdsson, S., et al. (2010). Genome-wide association studies of MRI-defined brain infarcts: meta-analysis from the CHARGE Consortium. Stroke 41, 210–217.

Elder, G. A., Friedrich, V. L., Jr, Kang, C., Bosco, P., Gourov, A., Tu, P. H., et al. (1998). Requirement of heavy neurofilament subunit in the development of axons with large calibers. J. Cell Biol. 143, 195–205.

Forest, M., Iturria-Medina, Y., Goldman, J. S., Kleinman, C. L., Lovato, A., Oros Klein, K., et al. (2017). Gene networks show associations with seed region connectivity. Hum. Brain Mapp. 38, 3126–3140.

Fox, M. D., Buckner, R. L., Liu, H., Chakravarty, M. M., Lozano, A. M., and Pascual-Leone, A. (2014). Resting-state networks link invasive and noninvasive brain stimulation across diverse psychiatric and neurological diseases. Proceedings of the National Academy of Sciences 111, E4367–E4375.

Fraher, J., and Dockery, P. (1998). A strong myelin thickness-axon size correlation emerges in developing nerves despite independent growth of both parameters. J. Anat. 193 (Pt 2), 195–201.

French, L., and Paus, T. (2015). A FreeSurfer view of the cortical transcriptome generated from the Allen Human Brain Atlas. Front. Neurosci. 9, 323.

French, L., Tan, P. P. C., and Pavlidis, P. (2011). Large-Scale Analysis of Gene Expression and Connectivity in the Rodent Brain: Insights through Data Integration. Front. Neuroinform. 5, 12.

Friede, R. L. (1972). Control of myelin formation by axon caliber (with a model of the control mechanism). J. Comp. Neurol. 144, 233–252.

Ganzetti, M., Wenderoth, N., and Mantini, D. (2015). Mapping pathological changes in brain structure by combining T1- and T2-weighted MR imaging data. Neuroradiology 57, 917–928.

García-Díaz, B., Riquelme, R., Varela-Nieto, I., Jiménez, A. J., de Diego, I., Gómez-Conde, A. I., et al. (2015). Loss of lysophosphatidic acid receptor LPA1 alters oligodendrocyte differentiation and myelination in the mouse cerebral cortex. Brain Struct. Funct. 220, 3701–3720.

Glasser, M. F., Coalson, T. S., Robinson, E. C., Hacker, C. D., Harwell, J., Yacoub, E., et al. (2016). A multi-modal parcellation of human cerebral cortex. Nature 536, 171–178.

Glasser, M. F., Goyal, M. S., Preuss, T. M., Raichle, M. E., and Van Essen, D. C. (2014). Trends and properties of human cerebral cortex: correlations with cortical myelin content. Neuroimage 93 Pt 2, 165–175.

Glasser, M. F., and Van Essen, D. C. (2011). Mapping human cortical areas in vivo based on myelin content as revealed by T1- and T2-weighted MRI. J. Neurosci. 31, 11597–11616.

Grier, M. D., Carson, R. P., and Lagrange, A. H. (2015). Of mothers and myelin: Aberrant myelination phenotypes in mouse model of Angelman syndrome are dependent on maternal and dietary influences. Behav. Brain Res. 291, 260–267.

Harting, I., Seitz, A., Rating, D., Sartor, K., Zschocke, J., Janssen, B., et al. (2009). Abnormal myelination in Angelman syndrome. Eur. J. Paediatr. Neurol. 13, 271–276.

Hawrylycz, M. J., Lein, E. S., Guillozet-Bongaarts, A. L., Shen, E. H., Ng, L., Miller, J. A., et al. (2012). An anatomically comprehensive atlas of the adult human brain transcriptome. Nature 489, 391–399.

Hawrylycz, M., Miller, J. A., Menon, V., Feng, D., Dolbeare, T., Guillozet-Bongaarts, A. L., et al. (2015). Canonical genetic signatures of the adult human brain. Nat. Neurosci. 18, 1832–1844.

Iwatani, J., Ishida, T., Donishi, T., Ukai, S., Shinosaki, K., Terada, M., et al. (2015). Use of T1-weighted/T2-weighted magnetic resonance ratio images to elucidate changes in the schizophrenic brain. Brain Behav. 5, e00399.

Kim, J.-W., Naidich, T. P., Ely, B. A., Yacoub, E., de Martino, F., Fowkes, M. E., et al. (2016). Human habenula segmentation using myelin content. Neuroimage 130, 145–156.

Koenig, S. H. (1995). Classes of hydration sites at protein-water interfaces: the source of contrast in magnetic resonance imaging. Biophys. J. 69, 593–603.

Krishnan, V., and Zeichner, S. L. (2004). Host cell gene expression during human immunodeficiency virus type 1 latency and reactivation and effects of targeting genes that are differentially expressed in viral latency. J. Virol. 78, 9458–9473.

Kugathasan, R., Collier, D. A., Haddow, L. J., El Bouzidi, K., Edwards, S. G., Cartledge, J. D., et al. (2017). Diffuse White Matter Signal Abnormalities on Magnetic Resonance Imaging Are Associated With Human Immunodeficiency Virus Type 1 Viral Escape in the Central Nervous System Among Patients With Neurological Symptoms. Clin. Infect. Dis. 64, 1059–1065.

Lerch, J. P., van der Kouwe, A. J. W., Raznahan, A., Paus, T., Johansen-Berg, H., Miller, K. L., et al. (2017). Studying neuroanatomy using MRI. Nat. Neurosci. 20, 314–326.

Li, X., Zhang, T., Liu, T., Lv, J., Hu, X., Guo, L., et al. (2016). A data-driven method to study brain structural connectivities via joint analysis of microarray data and dMRI data. in 2016 IEEE 13th International Symposium on Biomedical Imaging (ISBI), 829–832.

Løken-Amsrud, K. I., Myhr, K.-M., Bakke, S. J., Beiske, A. G., Bjerve, K. S., Bjørnarå, B. T., et al. (2013). Retinol levels are associated with magnetic resonance imaging outcomes in multiple sclerosis. Mult. Scler. 19, 451–457.

MacKay, A. L., and Laule, C. (2016). Magnetic Resonance of Myelin Water: An in vivo Marker for Myelin. BPL 2, 71–91.

Mahfouz, A., Huisman, S. M. H., Lelieveldt, B. P. F., and Reinders, M. J. T. (2017). Brain transcriptome atlases: a computational perspective. Brain Struct. Funct. 222, 1557–1580.

Mistry, M., and Pavlidis, P. (2010). A cross-laboratory comparison of expression profiling data from normal human postmortem brain. Neuroscience 167, 384–395.

Müller, C., Bauer, N. M., Schäfer, I., and White, R. (2013). Making myelin basic protein -from mRNA transport to localized translation. Front. Cell. Neurosci. 7, 169.

NCBI Resource Coordinators (2016). Database resources of the National Center for Biotechnology Information. Nucleic Acids Res. 44, D7–19.

Ogan Mancarci, B., Toker, L., Tripathy, S. J., Li, B., Rocco, B., Sibille, E., et al. (2017). Cross-laboratory analysis of brain cell type transcriptomes with applications to interpretation of bulk tissue data. bioRxiv, 089219. doi:10.1101/089219.

Peng, K., Xu, W., Zheng, J., Huang, K., Wang, H., Tong, J., et al. (2013). The Disease and Gene Annotations (DGA): an annotation resource for human disease. Nucleic Acids Res. 41, D553–60.

Portales-Casamar, E., Ch'ng, C., Lui, F., St-Georges, N., Zoubarev, A., Lai, A. Y., et al. (2013). Neurocarta: aggregating and sharing disease-gene relations for the neurosciences. BMC Genomics 14, 129.

Richiardi, J., Altmann, A., Milazzo, A.-C., Chang, C., Chakravarty, M. M., Banaschewski, T., et al. (2015). BRAIN NETWORKS. Correlated gene expression supports synchronous activity in brain networks. Science 348, 1241–1244.

Rizzo, G., Veronese, M., Expert, P., Turkheimer, F. E., and Bertoldo, A. (2016). MENGA: A New Comprehensive Tool for the Integration of Neuroimaging Data and the Allen Human Brain Transcriptome Atlas. PLoS One 11, e0148744.

Romme, I. A. C., de Reus, M. A., Ophoff, R. A., Kahn, R. S., and van den Heuvel, M. P. (2017). Connectome Disconnectivity and Cortical Gene Expression in Patients With Schizophrenia. Biol. Psychiatry 81, 495–502.

Sahraian, M. A., Radue, E.-W., Haller, S., and Kappos, L. (2010). Black holes in multiple sclerosis: definition, evolution, and clinical correlations. Acta Neurol. Scand. 122, 1–8.

Santo-Domingo, J., and Demaurex, N. (2012). Perspectives on: SGP symposium on mitochondrial physiology and medicine: the renaissance of mitochondrial pH. J. Gen. Physiol. 139, 415–423.

Schmierer, K., Wheeler-Kingshott, C. A. M., Tozer, D. J., Boulby, P. A., Parkes, H. G., Yousry, T. A., et al. (2008). Quantitative magnetic resonance of postmortem multiple sclerosis brain before and after fixation. Magn. Reson. Med. 59, 268–277.

Schmitz, K., Brunkhorst, R., de Bruin, N., Mayer, C. A., Häussler, A., Ferreiros, N., et al. (2017). Dysregulation of lysophosphatidic acids in multiple sclerosis and autoimmune encephalomyelitis. Acta Neuropathol Commun 5, 42.

Schubert, U., Ott, D. E., Chertova, E. N., Welker, R., Tessmer, U., Princiotta, M. F., et al. (2000). Proteasome inhibition interferes with gag polyprotein processing, release, and maturation of HIV-1 and HIV-2. Proc. Natl. Acad. Sci. U. S. A. 97, 13057–13062.

Seeger, M., Ferrell, K., Frank, R., and Dubiel, W. (1997). HIV-1 tat inhibits the 20 S proteasome and its 11 S regulator-mediated activation. J. Biol. Chem. 272, 8145–8148.

Shen, E. H., Overly, C. C., and Jones, A. R. (2012). The Allen Human Brain Atlas: comprehensive gene expression mapping of the human brain. Trends Neurosci. 35, 711–714.

Shimoyama, M., de Pons, J., Hayman, G. T., Laulederkind, S. J. F., Liu, W., Nigam, R., et al. (2015). The Rat Genome Database 2015: genomic, phenotypic and environmental variations and disease. Nucleic Acids Res. 43, D743–50.

Shin, J., French, L., Xu, T., Leonard, G., Perron, M., Pike, G. B., et al. (2017). Cell-Specific Gene-Expression Profiles and Cortical Thickness in the Human Brain. Cereb. Cortex, 1–11.

Shi, Y., Lan, F., Matson, C., Mulligan, P., Whetstine, J. R., Cole, P. A., et al. (2004). Histone demethylation mediated by the nuclear amine oxidase homolog LSD1. Cell 119, 941–953.

Singh, N., Vrontakis, M., Parkinson, F., and Chelikani, P. (2011). Functional bitter taste receptors are expressed in brain cells. Biochem. Biophys. Res. Commun. 406, 146–151.

Sprawls, P. (2000). Magnetic Resonance Imaging: Principles, Methods, and Techniques. Medical Physics Publishing.

Stedehouder, J., Couey, J. J., Brizee, D., Hosseini, B., Slotman, J. A., Dirven, C. M. F., et al. (2017). Fast-spiking Parvalbumin Interneurons are Frequently Myelinated in the Cerebral Cortex of Mice and Humans. Cereb. Cortex 27, 5001–5013.

Stüber, C., Morawski, M., Schäfer, A., Labadie, C., Wähnert, M., Leuze, C., et al. (2014). Myelin and iron concentration in the human brain: a quantitative study of MRI contrast. Neuroimage 93 Pt 1, 95–106.

Takasaki, C., Yamasaki, M., Uchigashima, M., Konno, K., Yanagawa, Y., and Watanabe, M. (2010). Cytochemical and cytological properties of perineuronal oligodendrocytes in the mouse cortex. Eur. J. Neurosci. 32, 1326–1336.

Tasic, B., Menon, V., Nguyen, T. N., Kim, T. K., Jarsky, T., Yao, Z., et al. (2016). Adult mouse cortical cell taxonomy revealed by single cell transcriptomics. Nat. Neurosci. 19, 335–346.

Thakurela, S., Garding, A., Jung, R. B., Müller, C., Goebbels, S., White, R., et al. (2016). The transcriptome of mouse central nervous system myelin. Sci. Rep. 6, 25828.

The UniProt Consortium (2017). UniProt: the universal protein knowledgebase. Nucleic Acids Res. 45, D158–D169.

van der Knaap, M. S., and Valk, J. (2011). Magnetic Resonance of Myelination and Myelin Disorders. Springer Berlin Heidelberg.

Vymazal, J., Hajek, M., Patronas, N., Giedd, J. N., Bulte, J. W., Baumgarner, C., et al. (1995). The quantitative relation between T1-weighted and T2-weighted MRI of normal gray matter and iron concentration. J. Magn. Reson. Imaging 5, 554–560.

Weiner, J., 3rd, and Domaszewska, T. (2016). tmod: an R package for general and multivariate enrichment analysis. PeerJ Preprints doi:10.7287/peerj.preprints.2420v1.

Yong-Hing, C. J., Obenaus, A., Stryker, R., Tong, K., and Sarty, G. E. (2005). Magnetic resonance imaging and mathematical modeling of progressive formalin fixation of the human brain. Magn. Reson. Med. 54, 324–332.

Zeisel, A., Muñoz-Manchado, A. B., Codeluppi, S., Lönnerberg, P., La Manno, G., Juréus, A., et al. (2015). Cell types in the mouse cortex and hippocampus revealed by single-cell RNA-seq. Science 347, 1138–1142.

Zeng, H., Shen, E. H., Hohmann, J. G., Oh, S. W., Bernard, A., Royall, J. J., et al. (2012). Large-scale cellular-resolution gene profiling in human neocortex reveals species-specific molecular signatures. Cell 149, 483–496.

